# Sodium Chloride Promotes the Growth of Bacterial Soil Isolate and Antimicrobial Activity of Polymixin B

**DOI:** 10.1101/2024.08.24.609488

**Authors:** Diwakar Kumar Singh

## Abstract

The decomposition of organic matter in the soil, soil salinity, and soil acidity are influenced by soil microbiology, which also controls the recycling and processing of nutrients in the soil. These factors affect soil fertility and ecological stability. This manuscript is focused on the isolation of seven gram-negative bacteria found in the agricultural soil of The Neotia University campus, West Bengal, India. These isolates have been found to be mesophilic based on their study of their growth profiles, which revealed that under the same incubation conditions, the isolates displayed rising development patterns between 24 and 120 hours. The presence of sodium and potassium chloride modulates the growth and development of bacterial strains (DNI1, DNI2, DNI3, DNI4, DNI5, DNI6, and DNI7) during laboratory set up. The sodium and potassium chloride composition of culture media that effectively regulates the development of bacterial isolates has been determined using the combinational method of salt treatment. The ideal physical factor required for the growth and development of microorganisms has been demonstrated by the pH and temperature titration during this study. The powerful antibiotics known as polymyxin B, which are nonribosomal lipopeptides produced by *Paenibacillus polymyxa*, are especially effective against Gram-negative bacteria. Because multidrug-resistant Gram-negative bacteria have few other choices for treating infections, the use of polymyxins in clinical settings has increased despite their toxicity in the past. This study offers an update on the most current findings about the bioactivity of soil isolates and their significant relationships to temperature, pH, salt, and polymixin B sensitivity. The toxicity impact may be reduced by enhancing polymixin B’s antibacterial activity with salt treatment in clinical research.

## 1. Introduction

Microorganisms found in soil are essential to its functions and characteristics. It consists of viruses, bacteria, fungi, archaea, and protozoa [1]. Microorganisms play a role in the decomposition of soil organic matter and the transformation of nutrients through processes like oxidation, nitrification, ammonification, nitrogen fixation, and others [2]. They can also store carbon and nutrients in their biomass, which is mineralised by remaining microbes after they die [3]. The growth of human society depends on both conventional and organic agriculture; nevertheless, the use of polluted water and improper application of organic chemical fertilisers can raise the microbial load which may be pathogenic or normal soil microbiota [4]. Recent advances in molecular and analytical methodologies have led to a greater understanding of these processes and more effective strategies for modifying them for various ecosystem services [5-7]. The movement of nutrients within and between the different biotic and abiotic pools in the soil environment is known as nutrient cycling [8, 9]. Most of the substances that microorganisms affect is essential for the growth of living beings because bacteria, archaea, and fungi recycling a variety of important nutrients. Microorganisms have a major role in the presence of phosphate (by phosphorus mineralisation), sulphate (via sulphur oxidation), and nitrate (via nitrification) in soils. Thus, in order to preserve a valued and productive soil system, bacteria are necessary. Changes in land use or soil cultivation are examples of soil environment disturbances that can disrupt soil microbial populations and give a negative impact on soil nutrient cycle [10]. Furthermore, microorganisms are required for the humic compounds, whereas stable forms of organic carbon are required for the soil to sequester organic carbon [11] In the past, challenge tests and multi-factorial designs have been used to model the kinetic parameters of microorganisms in a variety of factors, such as temperature, acidity, salts, concentration of anti-microbial [12, 13]. Salinization is one of the main causes of soil deterioration in the modern world. Salt reduces soil microbial activity and stunts plant growth due to osmotic stress and toxic ions. Soil microorganisms are essential to soil health because they convert organic matter into easily available nutrients through the mineralisation process. Maintaining a high level of soil microbial activity is therefore essential [14].

The salinity of soil influences the existence of soil microbial diversity via osmotic effect and specific ion effects. Microbes can accumulate osmolytes to adjust low osmotic potential; however, this consumes a lot of energy, which leads to decreased growth and activity [15-17]. The maximum growth rate of bioprotective bacteria *Lactococcus piscium* CNCM I-4031 and the *Bronchothrix thermosphacta* is influenced by temperature, pH, and NaCl concentration [12]. However, the precise function of growth components may be established by a logical setup in which potassium chloride can be used in lieu of sodium chloride to ensure their significance in growth, human health and food safety [18, 19]. The chloride is widely available on earth and may be found in both terrestrial and marine habitats, its significance for the physiology and development of bacterial cells has been demonstrated in several report [20]. Salt has a dominant effect on food safety and quality because of its capacity to inhibit dangerous bacteria and lengthen food shelf life. The literature has provided a clear definition of how potassium chloride (KCl) and sodium chloride (NaCl) differ in their effects on the growth and acidity of lactic acid bacteria [20]. Antimicrobial activity of various carboxylic acid salts in sodium and potassium against a few prevalent foodborne pathogens and microorganisms linked to spoiling [21]. Sodium chloride is one of the abiotic stressors that affects the growth pattern and virulence-associated phenotypes of *E. coli* by activating certain pathways or modifying gene expression insight the cell [22]. High salt also protect the bacterial cell from several abiotic stresses including antibiotics. Some of report exemplified the mechanism of bacterial survival against antibiotics in presence of salt through increasing efflux pump expression [23]. The effects of root-associated *Paenibacillus polymyxa* groups on pepper plant growth promotion and induced systemic resistance. *P. polymyxa* is associated with rhizospheres of various crops including roots of pepper which promote the plant growth and systemic resistance to bacteria, fungus and other parasites [24]. It secreated exopolysecchride and polymixin antibiotics to support the systemic response of plant. *Acinetobacter baumannii* that is multidrug resistant has demonstrated a notable response to Polymyxin B and Rifampin in combination with Ampicillin or Cefoperazone in an in vitro investigation [25].

*Salmonella typhimurium* NCCP10812 and *Salmonella enteritidis* NCCP12243 may have their heat resistance, antibiotic susceptibility, and Caco-2 cell invasion assessed in relation to NaCl [26]. Elevated sodium chloride concentrations boost ibuprofen’s microbicidal effect on common cystic fibrosis pathogens [27]. The activity of antimicrobial peptides (AMPs) can be enhanced by sodium chloride treatment in a *Staphylococcus aureus*, halotolerant bacteria [28]. Antibiotics called polymyxins are frequently used as a last resort to treat Gram-negative bacteria that can be fatal. However, renal toxicity caused by polymyxin is a crucial dose-limiting factor that might result in less-than-ideal therapy [29]. Since there are currently few other alternatives for treating infections caused by multidrug-resistant Gram-negative organisms, the therapeutic usage of polymyxins whose toxicity has previously limited their use is increasing. Concurrently, an extensive number of semisynthetic and synthetic polymyxin analogues have been created recently in an effort to lessen the nephrotoxicity of the natural compounds. Despite these initiatives, no clinically approved polymyxin analogues have been discovered to yet. But this could soon change because three new polymyxin analogues are now undergoing therapeutic testing [30, 31]. Strong activity of polymyxin B is linked to super-stoichiometric accumulation that lasts a long time and is mediated by weak binding to lipid A of gram negative bacteria [32].

## 2. MATERIALS AND METHODS

### 2.1. Isolation of bacteria from agriculture soil

Isolation of bacteria from rice field of agriculture soil of The Neotia University, West Bengal, India. The soil samples were collected and brought to the lab in sterilised zip-lock polythene bags. The samples were stored at 4 °C in a refrigerator for further processing. In order to isolate bacteria from soil samples, 0.5g of the sample was reconstituted in 10ml of sterile distilled water using the conventional serial dilution plate method. The samples were then serially diluted 10^−4^ to 10^−8^ times. The solid plate was prepared by adding 2% (w/v) of agar in broth before autoclaving and used for plate preparation during aseptic condition in the laboratory.

A 100 μL aliquot of 10^−8^ was used to spread on a sterile mannitol agar plate containing 3% sodium chloride. The plate was then incubated for 72 hours at 37°C. We observed seven dominant colonies (DNI 1, DNI 2, DNI 3, DNI 4, DNI 5, DNI 6, DNI 7) were grown on the plates, and each colony’s features were used for laboratory examination under the optimum condition. The separate colonies were maintained on an agar plate at 4 °C, Glycerol stock at - 80°C for biochemical characterisation in order to facilitate subsequent identification.

### 2.2. Molecular identification of the bacterial isolates

The bacterial isolate DNI1, DNI2, DNI3, DNI4, DNI5, DNI6, DNI7, DNI8 were identified using 16S rDNA sequencing. The bacterial universal primers, i.e., 5′-AGAGTTTGATCCTGGCTCAG-3′ primers (27F) and 5′-GGTTACCTTGTTACGACTT-3′ primers (1492R) were used to amplify the 16S rRNA gene.

### 2.3. Effect of NaCl & KCl concentrations on growth of bacterial isolates

The seven colonies of isolated bacteria were streaked over the mannitol agar plate and inoculated separately in different test tube. Mannitol broth was prepared by using 1g of D-Mannitol and 1g of Peptone with 100 ml distilled water both in presence and absence of salts. Bacterial isolates, 1:100 dilutions were cultured at 37°C for 24 to 48 hours after being inoculated in mannitol broth with varying salt concentrations (2%, 5%, or a mixture of salts containing different combination). The differential existence or lack of development in the culture tubes was observed and noted for the treatment analysis. The growth potential of the isolates at the specified salt concentration was determined by measuring the turbidity that developed in culture tubes using a spectrophotometer to 600 nM at different time frame (24h, 48h, 96h, and 120h)

### 2.4. Exposure of bacterial isolates to different pH

The bacterial isolates were inoculated in mannitol broth, 1:100 dilutions with pH of 6 and 7 and incubated at 37°C for 96h. The bacterial isolates exhibited a distinct growth pattern at varying pH levels, with the maximum growth occurring at pH = 7 when examined the growth pattern by spectrophotometer, O. D=600nM. Thus, the ideal pH =7.4 was used throughout the experiment to ascertain the varied responses of the isolates.

### 2.5. Effect of Temperature on growth of isolates

The mannitol broth was inoculated with bacterial isolates, 1:100 at 20°C, 27°C, 37°C, and 45°C to determine their sensitivity within the mesophilic range of temperature. A 1% suspected bacterial isolate was added to 10 mL of mannitol broth tubes for inoculation. A UV-VIS spectrophotometer was used to track the development, and measurements were made at O.D 600 nm to ascertain their suitability for activity.

## Result

### 1. Growth of bacterial isolates in Mannitol broth

The seven isolated colonies were streaked over the mannitol agar plate and thereafter utilised for individual inoculation in distinct mannitol broths for further analysis. The culture tubes were placed at 37 °C for differential analysis of growth of bacterial isolates at O.D 600nM and CFU counting. In 2.5% sodium chloride, the isolates grow more rapidly, but in the same concentration of potassium chloride, they are inhibited (Fig. 1.A). The findings may be confirmed with alternative salt combinations, and it was shown that 1.5% NaCl and 0.5% KCl are equally useful combinations for promoting the growth of isolated bacteria, as shown in Fig. 1.A. To investigate how bacterial isolates, behave in additional research with increasing salt concentrations. The isolates respond to growth more strongly in the presence of 5% KCl, whereas 5% NaCl suppresses growth. When compared to other salt combinations, 3.5% of KCl and 1.5% of NaCl show higher development, as seen in Fig. 1. B. The bacterial response is better at 2.5% NaCl, a lower salt concentration, than at higher salt concentrations as illustrated in Fig. 1C. The turbidity of the broth culture that has also been noted as illustrated in Fig. 1. D. The microbial diversity of the soil and other biotic and abiotic factors in agricultural soil may be the cause of the isolates’ varying responses.

**Table 1.**

| Table 1 |  |  |  |  |
| --- | --- | --- | --- | --- |
| Bacterial isolates | 5% KCl | 5%NaCl | 2.5% KCl + 2.5% NaCl | 3.75% KCl+ 1.5% NaCl |
| DNI 1 | + | -- | + | +++ |
| DNI 2 | ++ | -- | + | +++ |
| DNI 3 | + | -- | + | +++ |
| DNI 4 | +++ | -- | + | +++ |
| DNI 5 | ++ | -- | + | +++ |
| DNI 6 | +++ | -- | + | +++ |
| DNI 7 | +++ | -- | + | +++ |

**Table 2.**

| Table 2 |  |  |  |  |
| --- | --- | --- | --- | --- |
| Bacterial isolates | 2.5% KCl | 2.5% NaCl | 1% KCl + 1% NaCl | 0.5% KCl+ 1.5% NaCl |
| DNI 1 | -- | +++ | + | ++ |
| DNI 2 | -- | +++ | + | ++ |
| DNI 3 | -- | +++ | + | ++ |
| DNI 4 | - | +++ | + | ++ |
| DNI 5 | - | +++ | + | ++ |
| DNI 6 | - | +++ | + | ++ |
| DNI 7 | - | +++ | + | ++ |

**Fig. 1.**
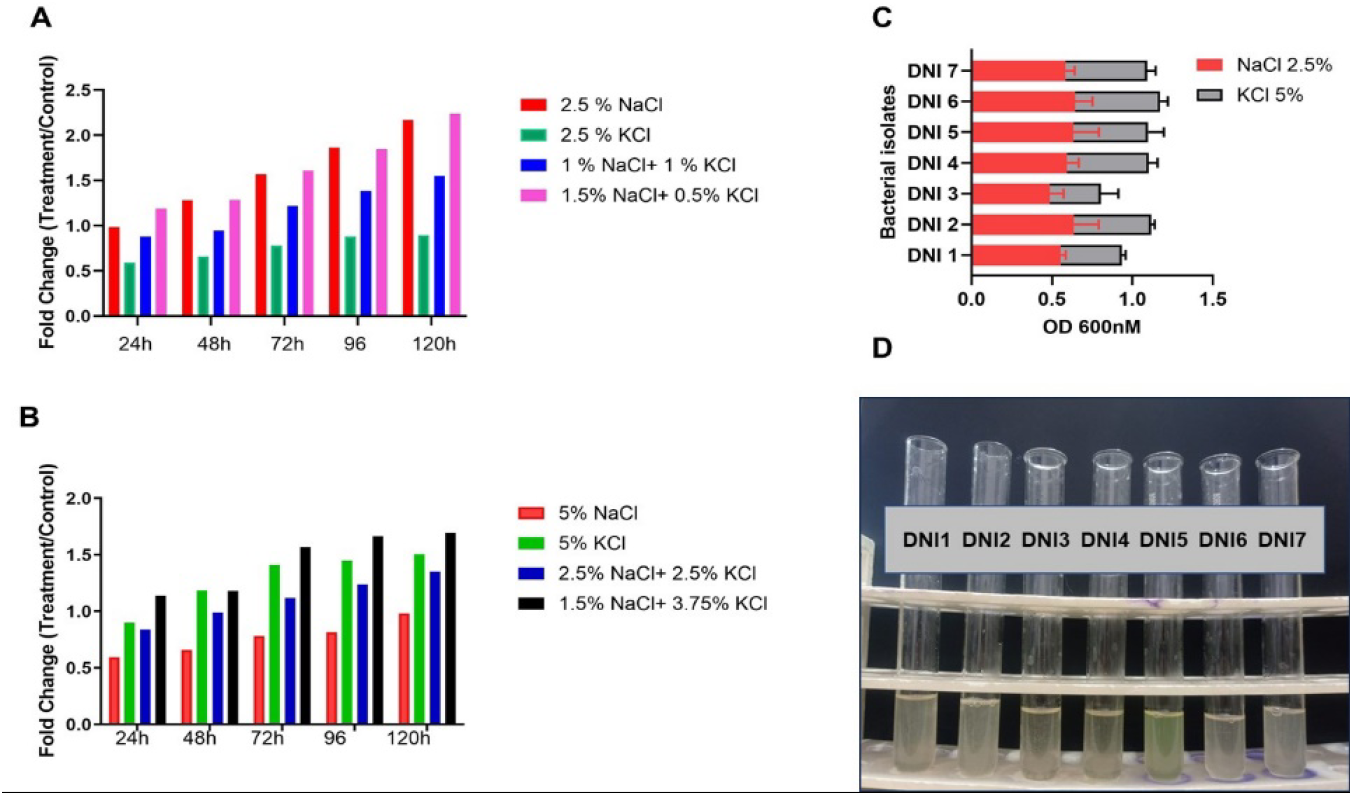
Growth pattern of bacterial isolates in presence of salt: The bacterial isolates from the soil grown in Mannitol broth at both higher and lower salt concentrations, monitored the growth pattern at O.D 600nM through spectrophotometer. A 2.5% treatment with sodium chloride exhibits more growth than 2.5% KCl and its other combination. B. Compared to 5% sodium chloride and its other combination, 5% potassium chloride exhibits greater growth. C. Compared to 5% potassium chloride, 2.5% sodium chloride is a more potent stimulant. DNI 2 bacterial isolate was used to investigate the sensitivity of both sodium chloride and potassium chloride as shown in A & B, D. The finding is strengthened and a similar growth trend is observed in the turbidity of the bacterial isolates, DNI 1, DNI 2, DNI 3, DNI 4, DNI 5, DNI 6, and DNI 7.

### 2. Effect of Temperature and pH on growth of microbes

Soil heterogeneity can be acquired (pedologically) or inherited (geologically), or it might be a combination of the two. The cultivation of soil has an effect on soil heterogeneity and microbial diversity. The bacterial isolates react differently when exposed to varying pH and temperature. After testing each isolate to determine the ideal physical condition needed for the growth and development of soil isolates, it was revealed that an acidic pH was not appropriate for bacterial growth when the growth pattern was examined at pH values of 6 and 7 (Fig. 2 B). Among the mesophilic temperature range, 37 °C is a more appropriate temperature, as shown in Figs. 2 A using O. D 600nM in heat map, The DNI2 is a study subject to be shown here. The other isolates exhibited a similar spectrum of bioactivity.

**Fig. 2.**
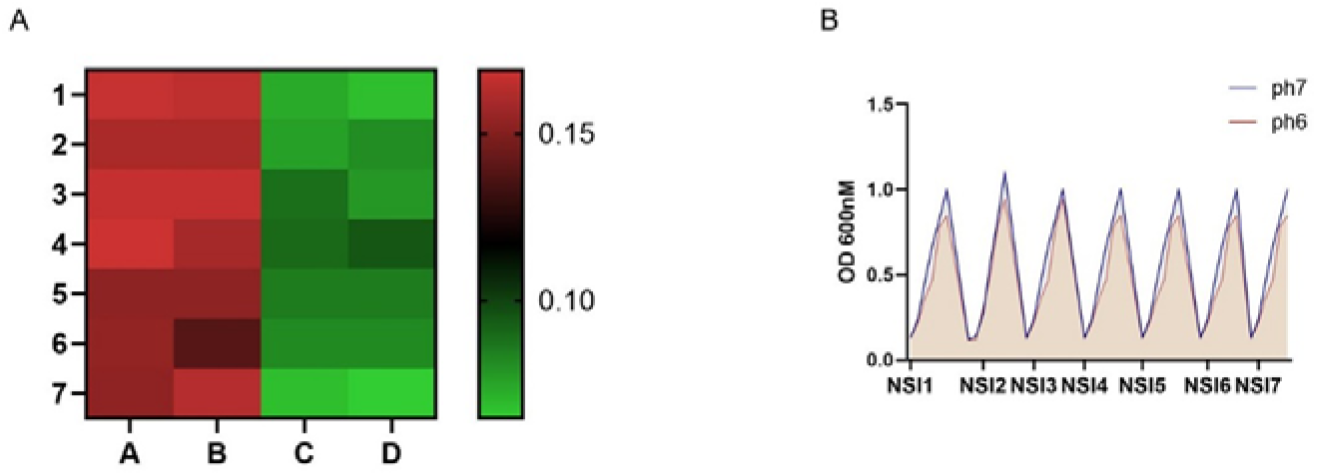
Growth pattern of bacterial isolates in presence differential physical factors. The soil-derived bacterial isolates grown under mesophilic laboratory conditions in mannitol broth. Intermediate temperature range is suitable for growth of isolates, such as temperature 27 °C and 37 °C, improved their responsiveness. A. The ideal 37 °C for the growth and development of isolates. B. The pH of a bacterium is another important physical factor that may be used to assess its level of sensitivity. Seven bacterial isolates have shown comparably better growth conditions at pH=7. These parameters were monitored during the course of the 96-hour treatment.

### 3. Antimicrobial activity of Polymixin B

The polymyxin B is an antibacterial polypeptide that acts quickly to kill bacteria by dissolving the lipid components of membranes, such as phospholipids and lipopolysaccharide. The seven colonies were sensitive to polymixin B when tested in this study (Fig. 3 A). The optimum antimicrobial activity of the molecule was examined with mannitol broth and solid agar plate under the laboratory condition. 2μg/ml of polymixin B was used as an effective concentration separately and with combination of sodium chloride (1% & 2%) and observed that sodium chloride enhanced the activity of polymixin B against DNI 2 as shown in Fig. 3B, D & E. The polymixin B activity was also estimated using a polymixin B strip (HiMedia) and found 16μg/ml was an effective concentration of the treatment in mannitol agar plate as illustrated in Fig. The activity of polymixin B was enhance when activity tested on the nitrate agar plate in place of mannitol. The MIC of polymixin B against bacterial isolates was determine 2μg/ml using different concentration of 0.5μg/ml to 8μg/ml on the basis of optical density of culture by spectrophotometer (Fig. 3C). The drug effectiveness demonstrated by colony forming unit per milliliter after 24h of treatment to revalidate the biological data as shown in Fig. 3 E.

**Fig. 3.**
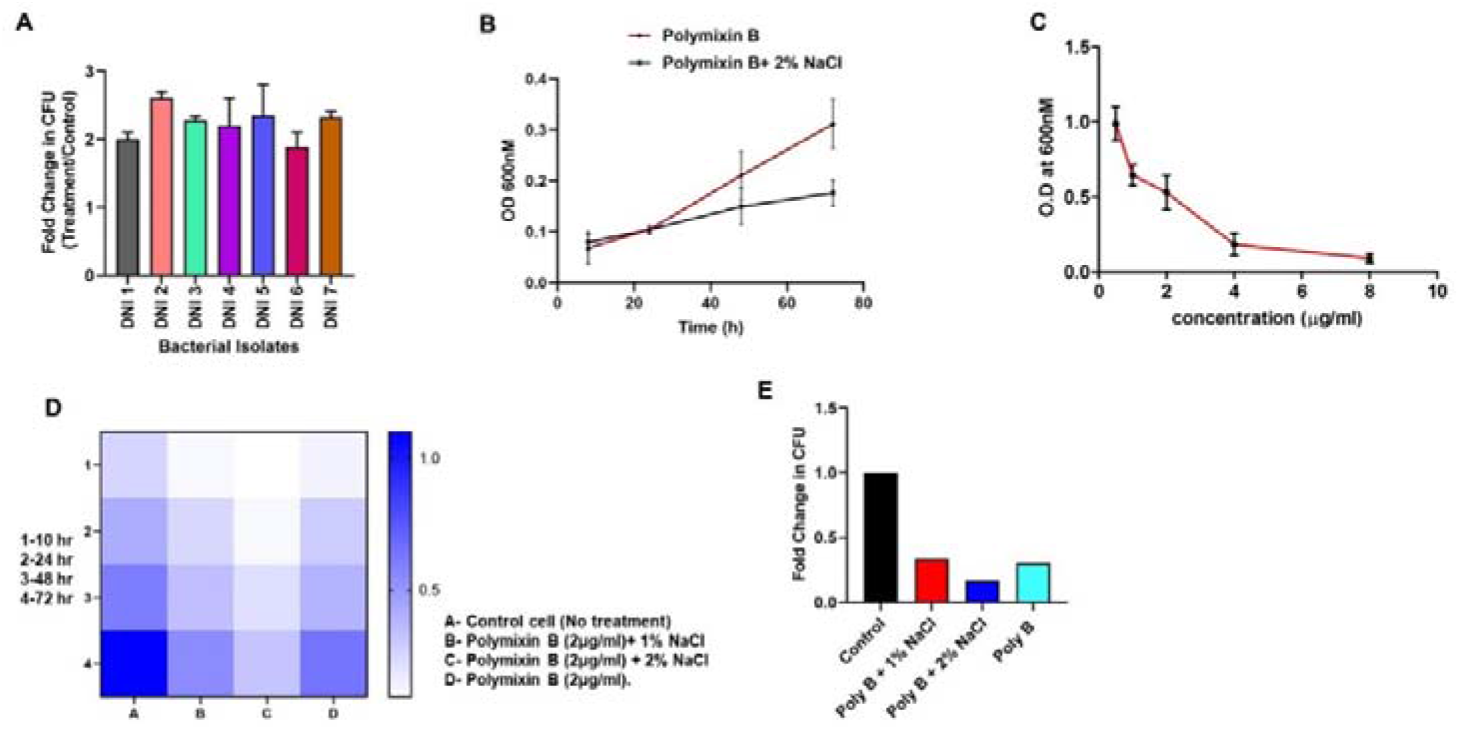
Antibacterial activity of Polymixin B: A. Seven bacterial isolates grown independently in mannitol broth exhibit distinct responses by treatment with 2μg/ml of Polymixin B; B. Polymixin B exhibits a significant increase in antibacterial activity with and without NaCl when treated for 48 and 72 hours on DNI 2 of Isolates. C. Polymixin B treatment of DNI 2 at concentrations ranging from 0.5μg/ml to 8μg/ml has demonstrated differential action and shows minimum inhibitory concentration (MIC) of polymixin B 2μg/ml. D. The bacterial Isolate DNI 2 was exposed to Polymixin B alone and in combination with NaCl for the duration of 10, 24, 48, and 72 hours and optical density is measured and represented here as a Heat map. E. Treatment of culture continue for 48h and live bacteria was enumerated by CFU through dilution platting.

## Discussion

The most often utilized potassium source is potassium chloride, and as a result of its constant usage, salt accumulation in plants and soil is increasing frequently. Overabundance of accessible ions can act as a biocide in the soil and induce a number of physiological problems in living things [33]. The potassium (K+) deficit caused by sodium is more common in salt-affected soils, which disturbs the K+/Na+ ratio in saline soil. However, this ratio needs to be maintained normal in order to preserve the balance of the soil and living things [34]. The sensitivity of the seven bacterial isolates (DNI 1, DNI 2, DNI 3, DNI 4, DNI 5, DNI 6, and DNI 7) to potassium and sodium chloride was tested using 2.5%, 1% and 1.5% of sodium chloride separately and with combination of potassium chloride (0%, 1% and 0.5%). 2.5% sodium chloride is more effective than other concentrations to promote the growth of bacterial isolates, and a combination treatment with 0.5% potassium chloride and 1.5% sodium chloride shows almost equivalent results (Fig. 1 A). When 2% potassium chloride was substituted for 2% sodium chloride, the results were reversed, and this lower range of salt therapy displays inhibitory effects (Fig. 1A). Numerous reports in the literature have substantiated sodium chloride’s bioactivity in analysis microbiome.

Thus, NaCl significantly influenced the protease activity, glucosidase activity, anaerobic fermentation, and microbial community distribution of soil [35]. Sodium and potassium chloride regulated the development and acid production of lactic acid bacteria [36]. Additionally, It modulated the expression of genes related to Bacillus sp. strain “SX4”‘s salt tolerance and rhamnolipid aggregation behaviour and antifungal activity [37, 38]. We can now be confident that sodium chloride, which is present in lower concentrations of salt, was a significant factor in the proliferation of the bacteria in laboratory set up. We broadened the scope of our research and looked into the potential role of higher concentration of salts (NaCl/KCl) in controlling bacterial growth. It observed that the same bacterial isolates grew more when exposed to 5% potassium chloride, but less when exposed to 5% sodium chloride. This function has been further confirmed by combining sodium chloride 2.5%, 1.5% with potassium chloride and it revealed that 1.5% NaCl and 3.75% KCl successfully promote bacterial growth when compared to other combinations in the laboratory (Fig. 1B). An extension of the study is subsequently examined the bioactivity of bacterial isolates in presence of 2.5% NaCl and 5% KCl, and the results indicated that lower salt concentration is more efficient in modulating development than higher concentration (Fig. 1C). In addition, Potassium chloride, Sodium lactate, and Sodium citrate reduced *Pseudomonas aeruginosa’s* antimicrobial resistance and virulence [39]. Potassium chloride effected the bioactivity of microbes that has been demonstrated using the salt replacement process[19, 40]. It also effected the growth, nitrogen mineralisation and soil microbial activity which is reported recently[21, 36]. The substitution of salt is illustrated to define the significance of NaCl and KCl on the growth and other bioactivity of *Listeria monocytogenes* [18]. The pH and temperature is a significant and important physical partners which regulates the bioactivity of soil isolates for maintaining the homeostasis. We observed, these bacterial isolates activity is decreased in acidic pH and low temperature while they retain the activity at higher temperature as shown in heat map assay (Fig.2 A) and neutral pH (Fig.2B). The bacterial isolates show optimum growth activity at 37 °C while moderate activity at 27 °C. This optimized physical parameters has helped us to determine the related bioactivity under the laboratory condition.

The microbial community in soil is regulated by the heterogeneous blend of biotic and abiotic components that make up the soil [41, 42]. We examined the antibacterial activity of polymixin B, which is generated by *Paenibacillus polymyxa*, against the seven bacterial isolates. In this investigation, we revealed that all isolates are susceptible to polymixinB (2μg/ml), a nonribosomal lipopeptide (Fig. 3A). In this investigation, DNI 2 is significantly more sensitive than other DNIs (Fig. 3A). When we examined the impact of sodium chloride on the isolated bacterium’s bioactivity. The antibacterial activity of polymixin B is increased when we treat bacterial isolate DNI 2 with 2μg/ml of the substance both the presence and absence of sodium chloride (Fig. 3 B). The concentration of polymixin B, which ranges from 0.5μg/ml to 8μg/ml, was used to determine the minimum inhibitory concentration (MIC) of the bacterial isolates. The corresponding optical density is shown in Fig. 3 C. As seen in Fig. 3 D, the antibacterial activity of polymixin B has been determined to be dependent on sodium chloride by titration of 2% and 1% sodium chloride across several time periods. The culture was cultivated for 48 hours, then slandered dilution platting was used to count the living colony. Fig. 3E shows the fold change in CFU as a function of the treatment.

The experiment was conducted three times, and each time a substantial difference was observed. Nevertheless, sodium chloride can boost the antimicrobial peptides’ (AMPs’) effectiveness against *Staphylococcus aureus* and enhance their microbicidal properties. Ibuprofen’s action against common pathogens that cause cystic fibrosis [27, 28].

Since diverse combinations of salts (NaCl & KCl) exhibited growth activity that is supportive for microbial isolates, these bacteria may be crucial for the maintenance and restoration of salt-impacted soils. The toxicity of polymixin B can be reduced by employing the same procedure, as these bacteria exhibited distinct antimicrobial activity in presence of it. The behaviour of microbes in the lab and utilise them as a model organism for titrating activity in the presence of novel biomolecules for human welfare, it is crucial to address the sodium & potassium chloride associated bioactivity of bacterial isolates through soil ecosystem which elaborated in this studies.

## Acknowledgments

The author expresses gratitude to The Neotia University for providing research support in the form of minor grants (TNU/R&D/M/10). Those that assisted in finishing the research project were Mr. Neelesh Pal, Ms. Tribeni Chowdhury, Ms. Puspa Gurung, and Ms. Soumi Kanji. The author was helped by Dr. Monisankar Bera in gathering soil samples from agricultural fields. The author is appreciative of this help.

